# The small RNA Teg16 represses *rsbV* and modulates SigB-dependent gene expression in *Staphylococcus aureus*

**DOI:** 10.64898/2026.06.01.729341

**Authors:** Sagar Pasham, Ananya Hota, Kaie Hall, Elise Turner, Sophie Lee, Olivia McGlaughlin, Karl M Thompson

## Abstract

*Staphylococcus aureus* relies on coordinated regulatory networks to adapt to environmental stress and host-associated conditions. The alternative sigma factor SigB plays a central role in this process and is controlled by the anti-anti-sigma factor RsbV, which functions as a key regulatory node in the pathway. While numerous small regulatory RNAs (sRNAs) have been identified in *S. aureus*, relatively few have been directly linked to the SigB stress response network. Here, we investigated the role of the small RNA Teg16 in post-transcriptional regulation of the SigB stress response pathway. Computational prediction identified a region of complementarity between Teg16 and the translational initiation region of *rsbV*. To test a potential regulatory effect based on this prediction, *teg16* was overexpressed, and *rsbV* transcript levels were measured by quantitative RT-PCR. Teg16 overexpression resulted in reduced *rsbV* transcript levels and decreased expression of SigB-dependent genes, including *asp23* and the carotenoid (*crt*) biosynthesis operon responsible for staphyloxanthin pigment production. In addition, strains carrying the *teg16* expression construct exhibited altered hemolytic activity under the conditions tested, suggesting effects on virulence-associated phenotypes. We further examined whether Teg16 influences the global regulator CodY and observed reduced *codY* transcript levels at early time points following *teg16* overexpression. Together, these results extend a previously identified regulatory relationship between Teg16 and CodY and raise the possibility of a feedback relationship linking post-transcriptional regulation to metabolic control. These findings identify Teg16 as a previously uncharacterized regulator that connects small RNA-mediated control to the SigB stress response network in *S. aureus*.

**Importance:** The alternative sigma factor SigB is a central regulator of stress adaptation in *Staphylococcus aureus* and influences both metabolism and virulence-associated phenotypes. While numerous small regulatory RNAs (sRNAs) have been identified in this organism, few have been functionally linked to control of the SigB pathway. Here, we identify the small RNA Teg16 as a regulator of *rsbV*, a key modulator of SigB activity. Teg16-dependent repression of *rsbV* is associated with reduced expression of SigB-dependent genes and measurable changes in phenotype, including decreased pigment production and altered hemolytic activity. In addition, our findings extend a previously identified relationship between Teg16 and the global regulator CodY, suggesting integration of post-transcriptional regulation with metabolic control. These results establish Teg16 as a previously uncharacterized component of the SigB regulatory network and provide new insight into how small RNAs contribute to stress adaptation in *S. aureus*.

## Introduction

*Staphylococcus aureus* is a versatile Gram-positive bacterium that functions both as a commensal colonizer and an opportunistic pathogen (1). As a colonizer, *S. aureus* persists on epithelial surfaces where nutrient availability, microbial competition, and host-derived stresses can fluctuate substantially. During infection, the organism encounters additional pressures, including oxidative stress, immune surveillance, antimicrobial exposure, and changes in tissue microenvironment (2). Survival under these conditions requires regulatory systems that allow the cell to rapidly adjust gene expression in response to environmental and metabolic cues. Thus, stress-response regulation is central to *S. aureus* persistence, adaptation, and virulence-associated physiology.

Among the regulatory systems that contribute to this adaptability is the alternative sigma factor SigB, a homologue of the stationary phase/general stress sigma factor in the firmicute model organism *Bacillus subtilis* (3–6). SigB plays a central role in mediating the general stress response in *S. aureus* (4, 5, 7–9) and influences a broad range of physiological processes, including stress resistance, metabolism, and virulence-associated phenotypes (4, 7, 8, 10, 11). SigB activity is tightly controlled through regulated interaction with the anti-sigma factor RsbW (5, 12, 13). The *sigB* operon encodes the regulatory proteins RsbU, RsbV, and RsbW, which are homologous to a similar genetic locus in *Bacillus subtilis* (5, 14–17). Under non-stress conditions, RsbW binds and sequesters SigB, preventing transcriptional activation (16, 18). Upon pathway activation, dephosphorylated RsbV binds RsbW, releasing SigB and enabling transcription of SigB-dependent genes (19, 20). This partner-switching mechanism places RsbV at a critical control point for modulating SigB activity. RsbU functions upstream in this pathway by controlling the phosphorylation state of RsbV (21).

The SigB regulon comprises a large set of genes involved in stress adaptation, metabolism, and virulence, and has been extensively characterized (22–25). SigB-dependent regulation contributes to environmental stress response and adaptation to host-associated conditions, linking stress response to persistence and pathogenesis (10, 11, 26, 27). The SigB regulon includes genes such as *asp23*, encoding alkaline shock protein 23 (28), and the *crt* operon involved in carotenoid biosynthesis and staphyloxanthin production (29, 30). In addition to direct transcriptional control, SigB intersects with broader regulatory networks, including modulation of *sarA* and repression of the *agr* quorum-sensing system (8, 31–35). Because *agr* is a major regulator of extracellular virulence factors, including hemolytic activity, these interactions place SigB within a larger regulatory network controlling extracellular proteins, protease activity, and virulence-associated physiology.

In addition to transcriptional regulation, small RNA-mediated post-transcriptional mechanisms are increasingly recognized as important contributors to gene regulation in bacteria (36–38). Studies of mRNA turnover and transcriptome dynamics have highlighted the importance of RNA stability in shaping gene expression responses under stress conditions (37, 38). Small regulatory RNAs (sRNAs) can further modulate mRNA stability or translation, enabling rapid adjustment of gene expression in response to changing environmental conditions (39, 40). In several well-characterized systems, sRNAs function within broader regulatory circuits, including those controlled by alternative sigma factors, as exemplified by the RpoE – RybB pathway in Gram-negative bacteria (41). Genome-wide studies have identified hundreds of sRNAs in *Staphylococcus aureus*, yet the functions of most remain poorly defined (42–46). Functional characterization of select sRNAs, including IsrR and Teg41, has demonstrated roles in metabolic regulation and virulence, respectively, highlighting the potential impact of the many uncharacterized sRNAs in this organism (47–52). Emerging evidence suggests that sRNA-sigma factor regulatory circuits may be more widespread than currently appreciated but remain relatively underexplored in Gram-positive bacteria, including *S. aureus*. Accordingly, relatively few sRNAs have been directly linked to central stress response pathways mediated by alternative sigma factors, such as the SigB regulatory system.

The small RNA Teg16 has emerged as a candidate regulator within these networks. Previous work demonstrated that *teg16* expression is strongly influenced by the global metabolic regulator CodY, with Δ *codY* mutants exhibiting substantial increases in *teg16* levels (53). Despite this regulatory connection, the functional role of Teg16 and its downstream targets has not been defined. Here, we investigated whether Teg16 contributes to post-transcriptional regulation of stress response pathways in *Staphylococcus aureus*. Using a combination of computational prediction and experimental analysis, we identify *rsbV* as a candidate target of Teg16 and demonstrate that Teg16 represses *rsbV* expression and modulates SigB-dependent gene expression. In addition, our findings expand the regulatory relationship between Teg16 and CodY, suggesting that Teg16 may participate in feedback interactions linking post-transcriptional regulation to metabolic control. Together, these results establish Teg16 as a previously uncharacterized regulator that connects small RNA-mediated control to the SigB stress response network in *S. aureus*.

## Materials and Methods

### Media and Growth Conditions

All *S. aureus* strains were grown in Tryptic Soy Broth (TSB) (KD Medical, Columbia, MD). Overnight cultures were grown in 5 mL of TSB in a 15 mL capped test tube in a Cel-Gro tissue culture rotator (Fisher Scientific, Pittsburgh, PA, USA), supplemented with chloramphenicol (TSB-Cm) as needed for plasmid selection to a final concentration of 10 μg / mL. For expression studies and downstream assessment of regulatory phenotypes, overnight cultures were diluted 1:500 in 25 mL of fresh TSB-Cm in a sterile 125 mL or 250 mL beveled glass Erlenmeyer flask (Fisher Scientific) in a WS27 shaking water bath (Shellab, Cornelius, OR) at 37°C at 150-160 rpm. For induction of plasmid-based *teg16* small RNA expression (Teg16 over-expression), TSB or TSB-Cm cultures were grown to an OD_600_ of 0.3 and supplemented with xylose to a final concentration of 2% and further incubated for an additional 30 to 180 minutes and a comparable strain with an empty vector (Vector Control). For the qualitative analysis of pigment, to streak for single colonies, or to create lawns for downstream techniques, cells were grown in Tryptic Soy Agar (TSA) (BD Biosciences, San Jose, CA, USA) supplemented with chloramphenicol (TSA-Cm) and xylose (TSA-Cm-Xyl) to final concentrations of 10 μg / mL and 2%, respectively for plasmid selection and expression induction as needed. Blood agar plates, used for hemolysis phenotyping, were made using TSA supplemented with 5% defibrinated sheep blood. Blood agar plates were supplemented with chloramphenicol to final a concentration of 10 μg / mL to select for plasmids.

All *Escherichia coli* strains were grown using LB (Lennox) Broth (KD Medical, Columbia, MD) or SOC Media (New England Biolabs, Ipswich, MA, USA), supplemented with ampicillin (LB-Amp) as needed for plasmid selection to a final concentration of 100 μg / mL. LB (Lennox) Agar plates, supplemented with ampicillin to a final concentration of 100 μg / mL, were used for selection of plasmid transformants for cloning reactions.

### Strains, Plasmids, Oligonucleotides, and Synthetic DNA

All *S. aureus* strains used in this study are derivatives of laboratory strains SH1000 and RN4220 and are listed in Table 1. All *E. coli* strains used in this study were for the purpose of molecular cloning and are derivatives of commercially available cloning strain – NEB® 5-alpha (NEB®-5α) (New England Biolabs, Ipswitch, MA) and are also listed in Table 1. All plasmids used in this study are derivatives of *E. coli* – *S. aureus* shuttle expression vector pEPSA5 and are listed in Table 1. Plasmid pEPSA5 facilitates xylose-inducible expression of genes via a xylose-responsive regulatory protein XylR and xylose-inducible promoter upstream of the multiple cloning sites. All oligonucleotides used in this study were obtained from Integrated DNA Technologies (IDT, Inc, Coralville, IA, USA) and are listed in Table 2. Oligonucleotide were used for the purpose of molecular cloning, construct verification, and expression profiling by quantitative reverse transcription PCR (qRT-PCR).

**Table 1.**
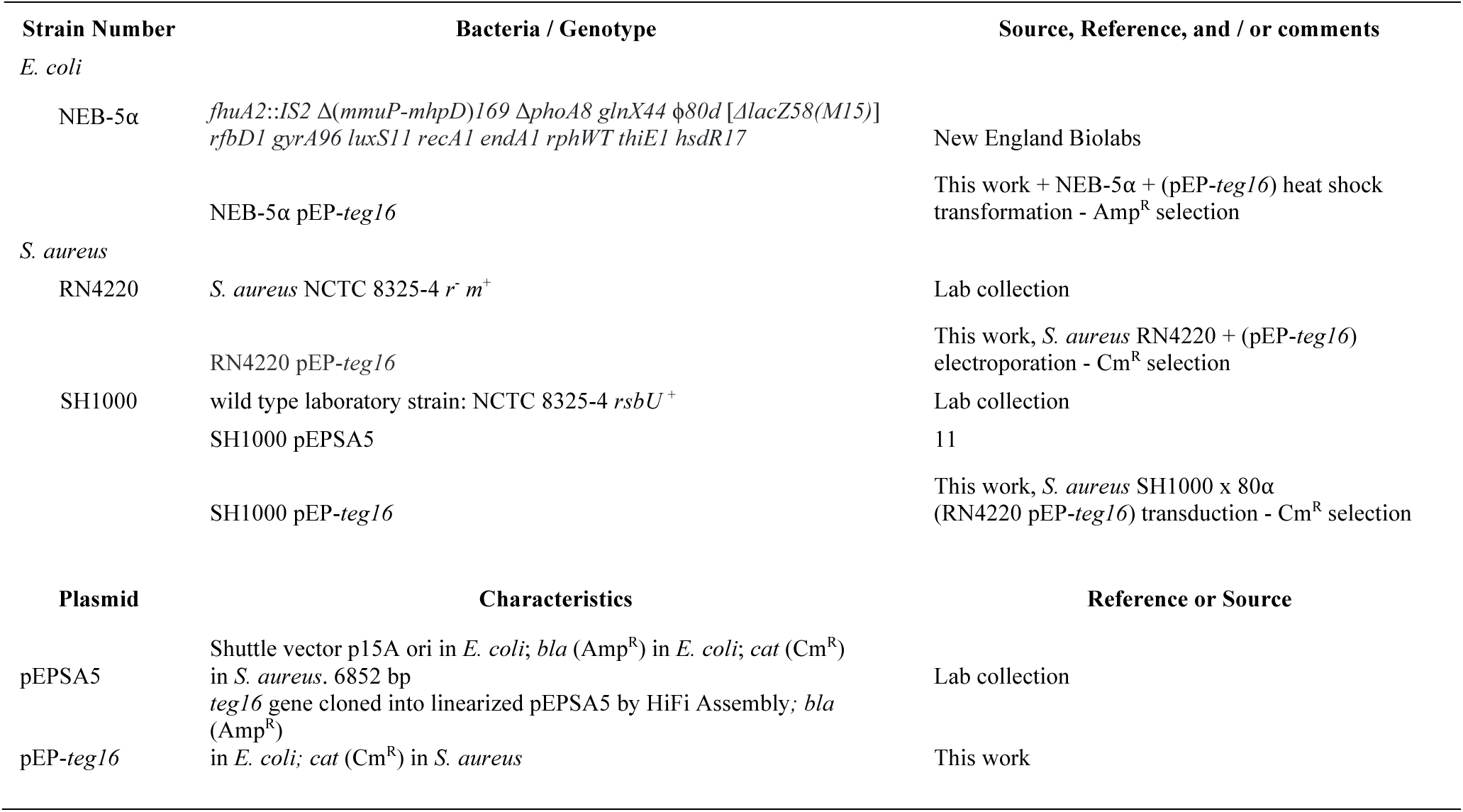
Strains and Plasmids.

**Table 2.**
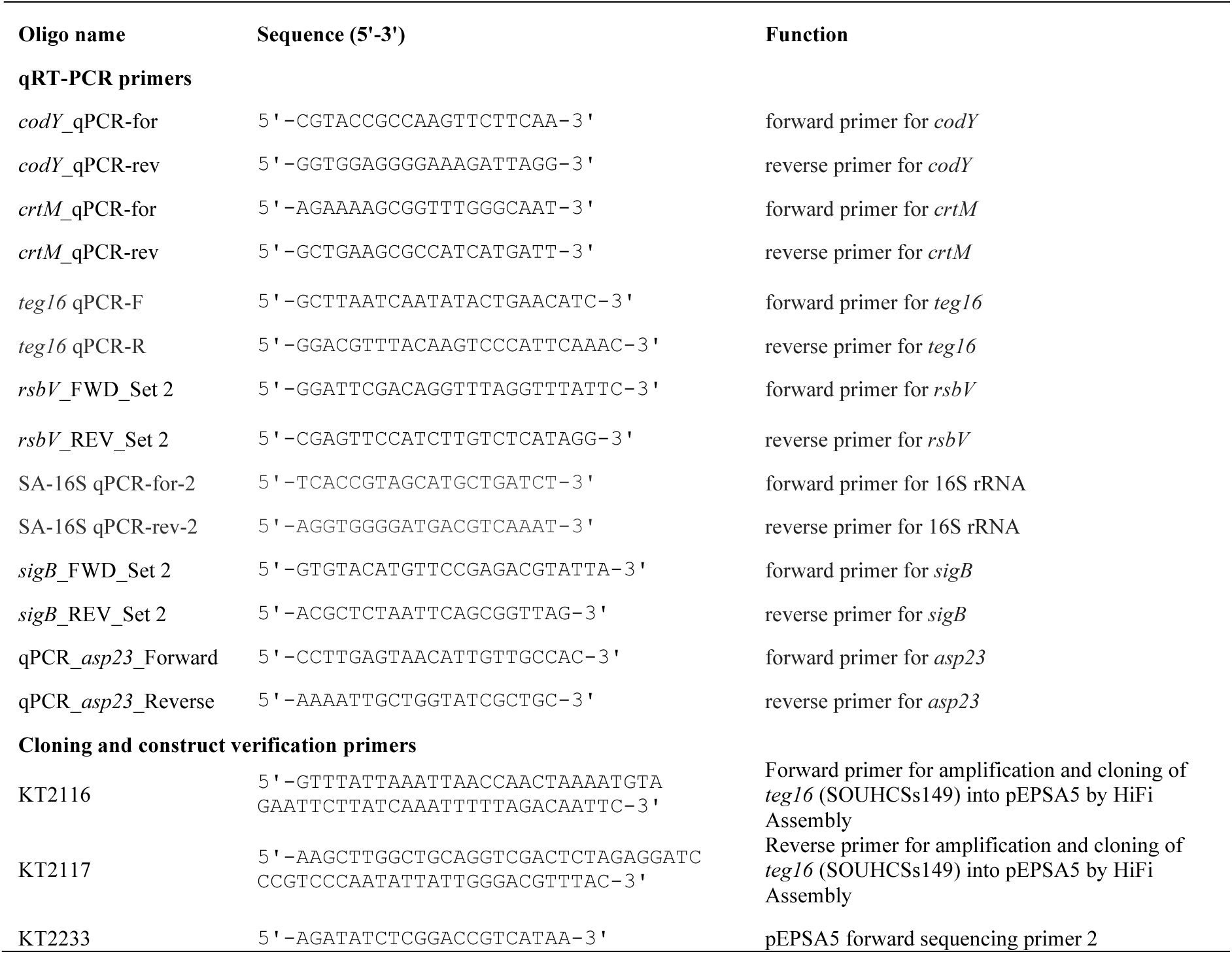
Synthetic Oligonucleotides used as primers in this study.

### Molecular Techniques and Genetic Engineering

Molecular techniques were performed using standard techniques. *S. aureus* genomic DNA from SH1000 was isolated, for molecular cloning, using the Column-Pure^TM^ Bacterial Genomic DNA Isolation Kit (Lamda Biotech, St. Louis, MO) according to manufacturer’s instructions, supplementing lysozyme for lysostaphin (Sigma Aldrich, St. Louis, MO) to a final concentration of 5 μg/mL. Plasmid DNA was isolated from *E. coli* or *S. aureus*, using the Column-Pure^TM^ Plasmid Mini-Prep Kit (Lamda Biotech, St. Louis, MO). PCR products were amplified using Red-Taq Master Mix (Lamda Biotech, St. Louis, MO). PCR products and plasmid restriction digests used for molecular cloning are purified using the Column-Pure^TM^ PCR Clean-Up Kit (Lamda Biotech, St. Louis, MO). Purified DNA and RNA were quantified using the Nanodrop^TM^ One/One^C^ Microvolume UV-Vis Spectrophotometer (Thermo Fisher Scientific, Waltham, MA, USA).

### Construction of pEPSA5-*teg16*

To create a xylose inducible allele of the small RNA *teg16*, the *teg16* gene was amplified from SH1000 genomic DNA using primers KT2116 and KT2117 (Table 2), creating 20-30 bp of homology to flanking regions of the pEPSA5 multiple cloning site (Table 1). PCR amplification was performed using the 2X RedTaq Master Mix (Lamda Biotech, St. Louis, MO) with an annealing temperature of 48°C and extension at 72°C for 30 seconds. The *teg16* PCR product was verified by agarose gel electrophoresis, purified, and quantified as described above. Purified pEPSA5 was subjected to restriction digestion using restriction endonucleases EcoRI and BamHI. The pEPSA5 EcoRI / BamHI digest was quantified and verified as described above. The purified pEPSA5 EcoRI / BamHI digest and *teg16* PCR product were recombined using the NEBuilder® HiFi Assembly Mastermix (New England Biolabs, Ipswitch, MA), according to manufacturer’s instructions with a 3:1 molar excess of insert to vector. An aliquot of the assembly reaction was mixed with an aliquot of NEB® 5-alpha (NEB®-5α) high efficiency chemically competent cells and subjected to heat shock, via a 30 second incubation at 42°C on a thermomixer (Thermo Fisher Scientific, Waltham, MA, USA). The transformation reaction was recovered by adding 500 μL of LB or SOC and incubated at 37°C at 200 rpm in a Thermomixer. A 100 μL aliquot of the recovery reaction was then spread on LB-Amp plates and incubated overnight at 37°C in a SI12 microbiological incubator (Shellab, OR). Transformants were transferred to another LB-Amp plate, to create a master plate, and screened by colony PCR for the presence of the *teg16* insert using oligonucleotide primers KT2233 (a vector specific forward primer) and KT2117 (an insert specific reverse primer). Plasmids from PCR positive colonies were purified and further verified by sanger sequencing. The resulting verified plasmid construct (pEPSA5-*teg16*, subsequently described as pEP-*teg16* or Teg16 Over-Expression in figures) was re-transformed into *E. coli* NEB®-5α for cryopreservation and electroporated into *S. aureus* RN4220 and subsequently transduced from RN4220 into SH1000.

### RNA Isolation

Total RNA was isolated using the FastRNA Pro Blue Kit (MP Biomedicals). Briefly, cell pellets from 1–10 mL cultures were harvested by centrifugation at 4000 rpm for 10 min using an Allegra X-14R refrigerated centrifuge (Beckman Coulter, Brea, CA, USA). Pellets were resuspended in 1 mL RNA Pro solution and transferred to lysing matrix B tubes for mechanical lysis. Samples were processed using a Precellys Dual-24 homogenizer (Bertin Instruments, Rockville, MD, USA) at 5000 rpm for 20 s for two cycles with a 5 min incubation on ice between cycles. Lysates were clarified by centrifugation, and RNA was extracted using chloroform and precipitated overnight at −80 °C with three volumes of ethanol. RNA pellets were collected by centrifugation at 14,000 rpm for 30 min at 4 °C, air-dried, and resuspended in nuclease-free water.

### Quantitative Reverse Transcriptase PCR (qRT-PCR)

Total RNA for qRT-PCR analysis was treated with the TURBO DNA-free™ Kit (Invitrogen, AM1907) to eliminate contaminating genomic DNA, followed by further purification using the Monarch® Spin RNA Cleanup Kit (NEB, T2040L). Purified, DNA-free RNA was then reverse transcribed into cDNA using the High-Capacity cDNA Reverse Transcription Kit (Applied Biosystems, 4368814). Quantitative PCR was carried out with PowerTrack™ SYBR Green Master Mix (Applied Biosystems, A46109) using a QIAGEN QIAquant 96 5plex Real-Time PCR system. Target-specific primers were used as listed in Table 2, with 16S rRNA serving as the internal reference gene, and non-template controls included to verify the absence of contamination. For each biological replicate, a minimum of two technical replicates was analyzed. Instrument-generated Ct values were used for the calculation of ΔCt and ΔΔCt, and relative expression levels were normalized to the vector control, which was set to 1.

### Qualitative assessment of *S. aureus* colony pigment

Pigmentation of *S. aureus* SH1000 strains carrying either the empty vector control pEPSA5 or pEPSA5-*teg16* was assessed qualitatively on solid medium and in liquid culture. For colony-based assessment, strains were transferred from cryostocks to TSA-Cm supplemented with chloramphenicol to a final concentration of 10 μg/mL and incubated at 37°C for 24 hours. No xylose was added to plates used for qualitative colony pigmentation assessment. Colony pigmentation was visually assessed based on yellow-orange intensity. For liquid-culture assessment, strains carrying either pEPSA5 or pEPSA5-*teg16* were grown in 125 mL of TSB-Cm to an OD600 of approximately 0.3. Cultures were then either left untreated or supplemented with xylose to a final concentration of 2% and incubated for an additional 30 minutes. Cells were harvested by centrifugation, and pellets were visually assessed for pigmentation. Pellets were then resuspended in 1 mL of ultrapure H2O and reassessed for yellow-orange pigmentation. Images shown are representative of at least three independent biological replicates.

### Methanol Extraction and Quantitative Analysis of Staphyloxanthin

Staphyloxanthin pigment was extracted with methanol and quantified using a protocol adapted from Campbell et al., with modifications (54). Briefly, vector control and *teg16 o*verexpression strains were grown in 25 mL of TSB-Cm in a shaking water bath at 37°C. Cultures were grown to an OD600 of approximately 0.3 and then either left untreated or supplemented with xylose to a final concentration of 2%. At 30 and 180 minutes after induction, 10 mL of culture was collected, and optical density was measured to monitor cell density. Cultures were centrifuged for 10 minutes at 4,000 rpm, and the supernatant was removed. Cell pellets were resuspended in 400 μL of methanol and incubated at 55°C for 30 minutes to extract pigment. Samples were then centrifuged for 5 minutes at 10,000 rpm, and 300 μL of the methanol extract was transferred to a clear 96-well plate in triplicate. Pigment levels were quantified by measuring absorbance at 465 nm using a microplate reader. Measurements were normalized to the corresponding vector control, which was set to a value of 1. At least three independent biological replicates were performed, and quantified data are presented as the mean ± SEM and analyzed for statistical significance.

### Qualitative assessment of *S. aureus* hemolysis

Hemolysis assays were performed on TSA plates supplemented with 5% defibrinated sheep blood and chloramphenicol to a final concentration of 10 μg/mL. Briefly, *S. aureus* SH1000 strains carrying either the empty vector control pEPSA5 or pEPSA5-*teg16* were restreaked from TSA-Cm plates and then transferred to blood agar plates prepared either without xylose or with xylose to a final concentration of 2%. Plates were incubated at 37°C for 24 and 48 hours. Hemolytic activity surrounding colonies was assessed visually based on zones of clearing. Representative images shown in Fig. 4 were obtained from plates without xylose, although similar qualitative hemolysis phenotypes were observed under inducing conditions. Images shown are representative of at least three independent biological replicates.

**Figure 1.**
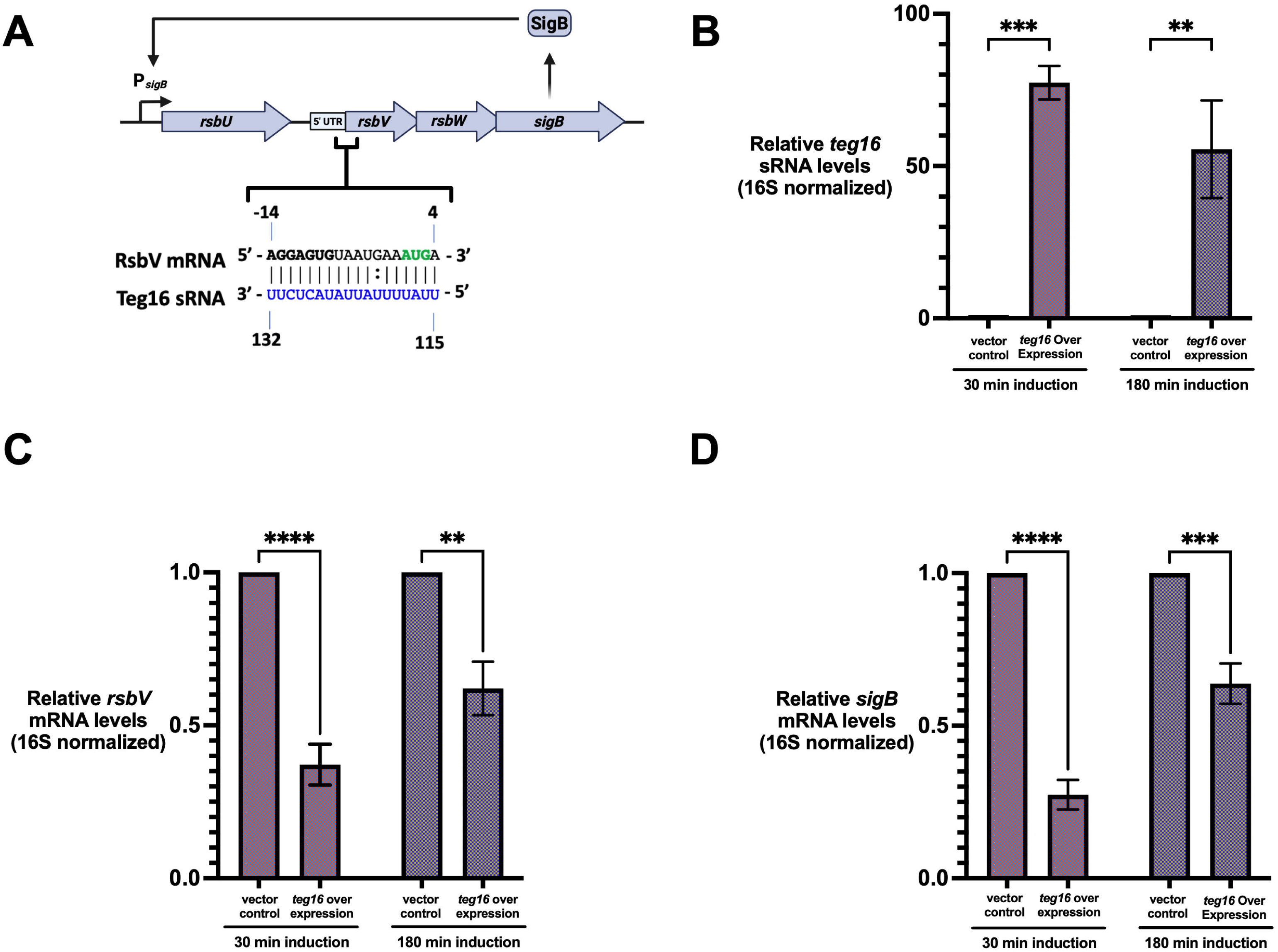
Teg16 is predicted to base-pair with the *rsbV* translational initiation region and represses *rsbV* and *sigB* expression. **(A)** Schematic representation of the *rsbU*-*rsbV*-*rsbW*-*sigB* operon and relative position of predicted base-pairing interaction between the small RNA Teg16 and the translational initiation region (TIR) of the *rsbV* mRNA. Nucleotides involved in base pairing are indicated. **(B)** Quantitative RT-PCR analysis confirming induction of *teg16* expression following addition of 2% xylose for 30 min or 180 min. Transcript levels were normalized to 16S rRNA and expressed relative to vector control. **(C-D)** Quantitative RT-PCR analysis of *rsbV* **(C)** and *sigB* **(D)** transcript levels following induction of *teg16* expression from a xylose-inducible promoter. Cultures were grown to mid-exponential phase (OD_600_ ∼ 0.3-0.5) and induced with 2% xylose for 30 min or 180 min prior to RNA isolation. Transcript levels were normalized to 16S rRNA and expressed relative to vector control. Data represent mean ± SEM from at least three independent biological replicates. Statistical significance was determined using two-way ANOVA with Šidák multiple comparisons test (\*\**P* < 0.01, \*\*\**P* < 0.001, \*\*\*\**P* < 0.0001), comparing *teg16* overexpression to vector control at each time point.

**Figure 2.**
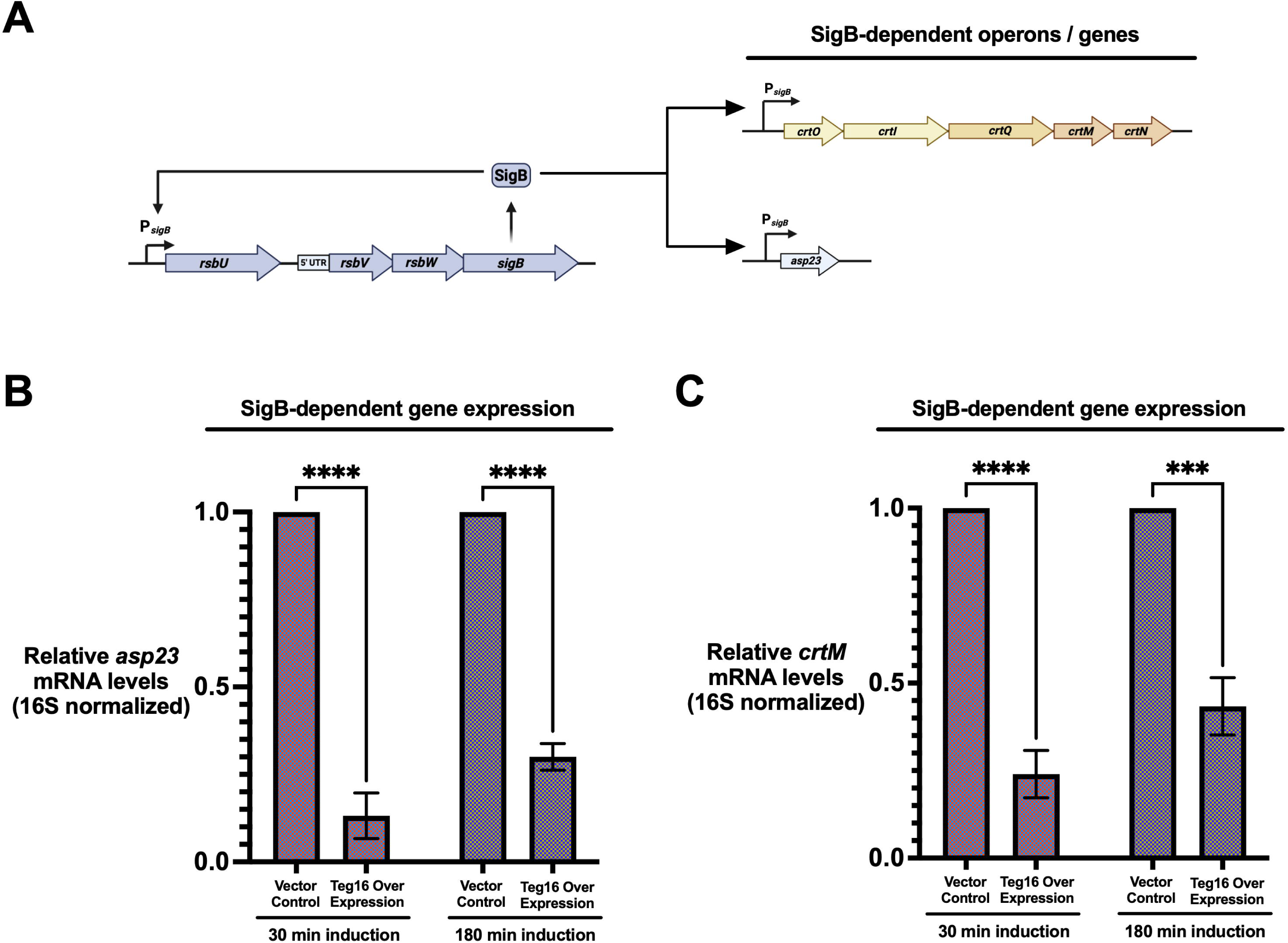
Teg16 represses SigB-dependent gene expression. **(A)** Schematic of the SigB regulatory pathway and downstream transcriptional outputs in *Staphylococcus aureus*, including the *rsbU*–*rsbV*–*rsbW*–*sigB* operon and SigB-dependent targets such as the *crt* operon and *asp23*. Quantitative RT-PCR analysis of SigB-regulated transcripts following induction of *teg16* overexpression. Expression levels of *asp23* **(B)** and *crtM* **(C)** mRNA were measured following induction with 2% xylose for 30 min or 180 min. Transcript levels were normalized to 16S rRNA and expressed relative to vector control. Data represent mean ± SEM from at least three independent biological replicates. Statistical significance was determined using two-way ANOVA with Sidak’s multiple comparisons test (\*\*\**P* < 0.001, \*\*\*\**P* < 0.0001).

**Figure 3.**
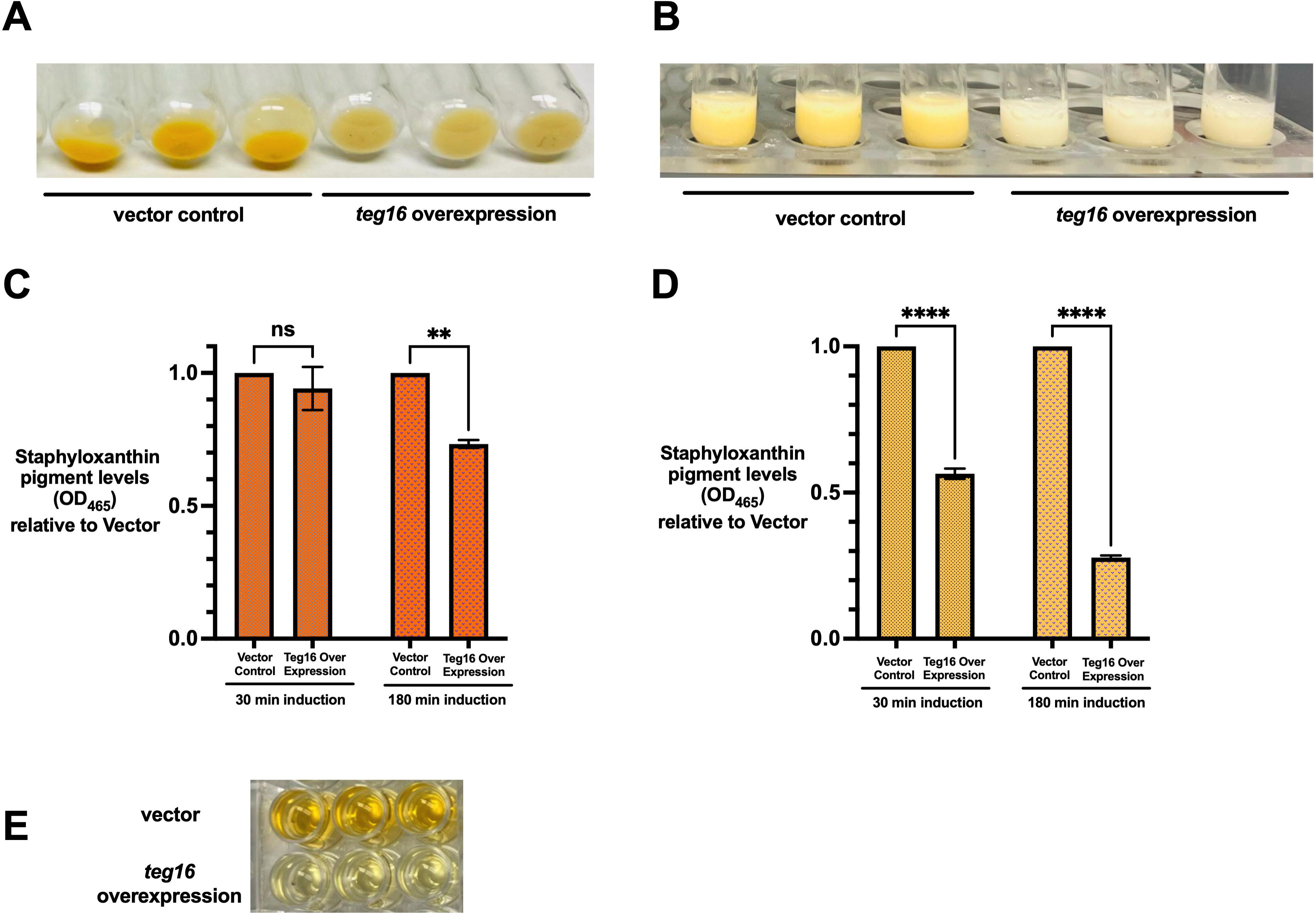
Teg16 overexpression represses staphyloxanthin production in *Staphylococcus aureus*. *Staphylococcus aureus* strains carrying either the empty vector control or inducible teg16 overexpression construct were grown under non-inducing and inducing conditions to assess effects on staphyloxanthin-associated pigmentation. **(A)** Representative cell pellets and **(B)** resuspended cell pellets from vector control and *teg16*-overexpressing cultures grown under non-inducing and inducing conditions. **(C, D)** Quantification of methanol-extracted pigment measured at OD465 from cultures grown without xylose **(C)** or with 2% xylose **(D)** at 30 and 180 minutes. Values were normalized to the corresponding vector control. **(E)** Representative resuspended methanol pigment extracts showing reduced pigment accumulation in *teg16*-overexpressing cultures. Data represent mean ± SEM from at least three independent biological replicates. Statistical significance was determined using two-way ANOVA followed by Šidák’s multiple-comparisons test (**P < 0.01, ****P < 0.0001).

**Figure 4.**
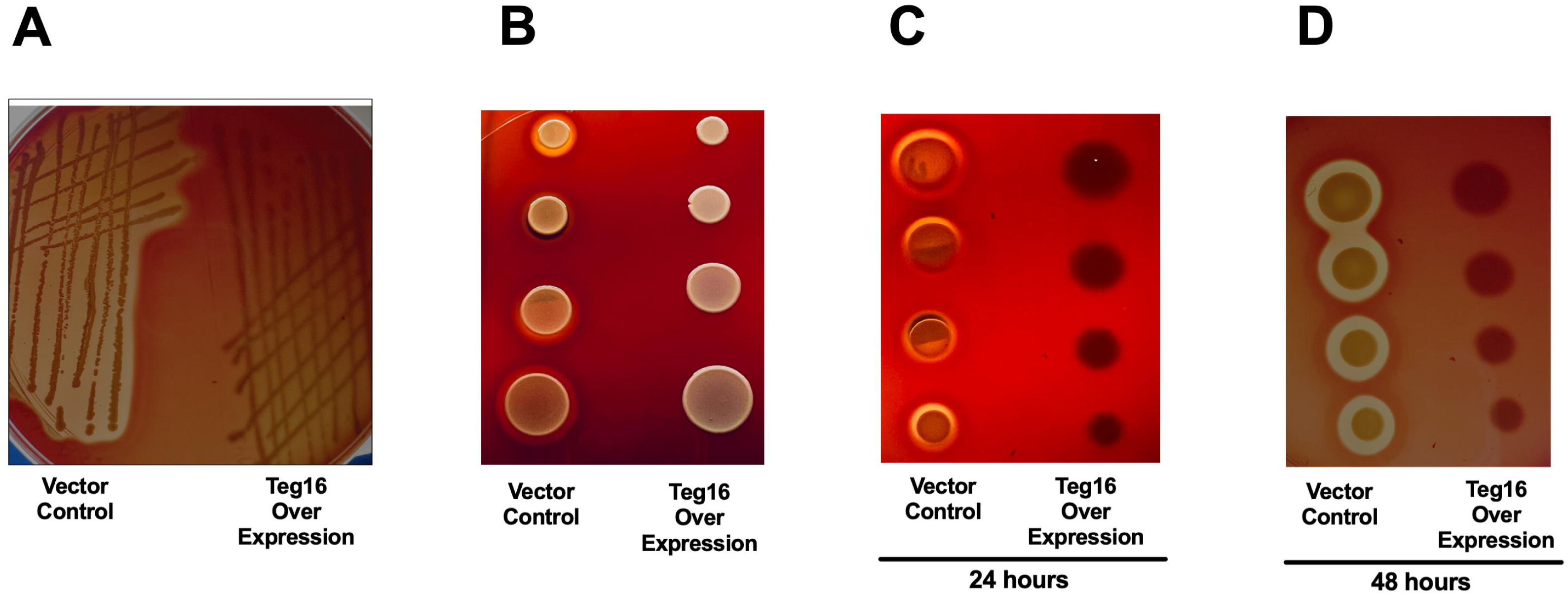
Teg16 alters hemolytic activity. To assess the effect of the *teg16* expression construct on hemolysis, *Staphylococcus aureus* strains carrying either the empty vector control pEPSA5 or pEPSA5-*teg16* were evaluated on blood agar plates. Representative images shown were obtained without xylose. **(A)** Representative streak plating on TSA supplemented with 5% defibrinated sheep blood showing differences in colony morphology and hemolytic activity between vector control and pEPSA5-*teg16* strains. **(B)** Spot assay, top view, demonstrating hemolytic activity following deposition of equal volumes of vector control and pEPSA5-*teg16* cultures onto blood agar plates. **(C–D)** Time-dependent hemolytic activity visualized from the underside of the plates at 24 hours **(C)** and 48 hours **(D)** post-incubation at 37°C. Strains carrying pEPSA5-*teg16* exhibited reduced zones of hemolysis compared to vector control. Images are representative of at least three independent biological replicates.

### Statistical Analysis

All quantitative experiments were performed using at least three independent biological replicates. Data are presented as the mean ± standard error of the mean (SEM). Statistical analyses were performed using GraphPad Prism (GraphPad Software, San Diego, CA, USA). Differences between groups were evaluated using two-way ANOVA with Šidák’s multiple comparisons test. Differences were considered statistically significant at *P* < 0.05. (\**P* < 0.05, \*\**P* < 0.01, \*\*\**P* < 0.001, \*\*\*\**P* < 0.0001).

## Results

### Teg16 represses the rsbV-rsbW-sigB operon in Staphylococcus aureus

To identify candidate targets of the small RNA Teg16, we performed in silico RNA–RNA interaction prediction. This analysis revealed a region of complementarity between Teg16 and the translational initiation region (TIR) of the *rsbV* mRNA, including sequences overlapping the ribosome binding site and start codon (Fig. 1A). The position of this predicted interaction suggests that Teg16 may repress *rsbV* expression by interfering with translation initiation and/or promoting transcript destabilization. To experimentally test this prediction, we examined the effect of Teg16 on *rsbV* expression using an inducible overexpression system. Quantitative RT-PCR analysis confirmed strong induction of *teg16* transcripts following xylose addition, with approximately 50-70-fold increased expression at both 30- and 180-minutes relative to vector control (Fig. 1B), demonstrating effective overexpression under the conditions tested. Under these conditions, *rsbV* transcript levels were reduced approximately 2.5-3.0-fold at 30 minutes and ∼2-fold at 180 minutes following Teg16 overexpression (Fig. 1C), consistent with negative post-transcriptional regulation. The magnitude and timing of this repression are consistent with a regulatory effect of Teg16 on *rsbV* expression. Because *rsbV*, *rsbW*, and *sigB* are co-transcribed as part of the SigB regulatory operon, we next assessed whether Teg16-mediated repression of *rsbV* is associated with altered *sigB* transcript levels. Consistent with this, *sigB* transcript levels were reduced approximately 3-fold at 30 minutes and ∼2-fold at 180 minutes following Teg16 induction (Fig. 1D), indicating that Teg16-dependent repression of the operon is associated with reduced expression of the SigB pathway. Together, these results support a model in which Teg16 represses the *rsbV*-*rsbW*-*sigB* operon, linking small RNA-mediated post-transcriptional control to modulation of the SigB stress response pathway in *Staphylococcus aureus*.

### Teg16 represses SigB-dependent gene expression

To determine whether Teg16-mediated repression of the *rsbV–rsbW–sigB* operon results in altered SigB activity, we examined expression of genes within the SigB regulon (Fig. 2A). The SigB-dependent gene *asp23*, a well-established marker of SigB activity, exhibited strong repression following Teg16 overexpression, with approximately 10-fold reduction at 30 minutes and ∼3-fold reduction at 180 minutes relative to vector control (Fig. 2B). This reduction is consistent with decreased SigB-dependent transcription. In addition, expression of *crtM*, a gene within the staphyloxanthin biosynthesis operon that is positively regulated by SigB, was also reduced following Teg16 induction, with approximately 4-fold repression at 30 minutes and ∼2.5-fold repression at 180 minutes (Fig. 2C). Because the *crt* operon contributes to carotenoid pigment production, these transcriptional changes suggest that Teg16-dependent modulation of the SigB pathway may have downstream phenotypic consequences. Notably, repression of both *asp23* and *crtM* was more pronounced at the early (30 minute) time point compared to 180 minutes, a pattern that parallels the temporal dynamics observed for *rsbV* and *sigB* repression (Figs. 1C & 1D). This consistency supports a model in which Teg16 exerts an early regulatory effect on the SigB pathway that may be partially attenuated over time, potentially reflecting compensatory regulatory responses. Together, these results indicate that Teg16-mediated repression of the *rsbV-rsbW-sigB* operon leads to reduced expression of SigB-dependent genes, consistent with decreased SigB pathway activity.

### Teg16 represses staphyloxanthin production

Because the *crt* operon contributes to carotenoid pigment production, we next assessed whether Teg16-mediated repression of the *rsbV*-*rsbW*-*sigB* operon was associated with altered pigmentation. Staphyloxanthin biosynthesis, which contributes to the characteristic yellow-orange pigmentation of *Staphylococcus aureus*, is positively regulated by SigB (Fig. 2A). To test this, we examined staphyloxanthin production following Teg16 overexpression under non-inducing (- xylose) and inducing (+ xylose) conditions at 30 and 180 minutes. Teg16-overexpressing strains exhibited visibly reduced pigmentation compared to vector control strains, with the reduction most apparent under inducing conditions, as observed in both cell pellets and resuspended cell pellets (Fig. 3A and 3B).

Consistent with these observations, spectrophotometric quantification of methanol-extracted pigment at OD_465_ demonstrated reduced staphyloxanthin accumulation in Teg16-overexpressing cultures grown without xylose at both 30 and 180 minutes (Fig. 3C). Following induction with 2% xylose, Teg16 overexpression resulted in a more pronounced reduction in staphyloxanthin levels, with approximately 2-fold reduced pigment accumulation at 30 minutes and approximately 3-fold reduced pigment accumulation at 180 minutes relative to vector control (Fig. 3D). Representative resuspended pigment extracts further illustrated reduced pigment accumulation in Teg16-overexpressing cultures (Fig. 3E). Together, these results demonstrate that Teg16 represses staphyloxanthin production in an induction-dependent manner, providing a phenotypic readout consistent with reduced SigB pathway activity.

### Teg16 alters hemolytic activity

Having established that Teg16 overexpression reduces a SigB-associated pigmentation phenotype, we next asked whether the *teg16* expression construct was also associated with altered hemolytic activity. Hemolytic activity is controlled by complex regulatory networks in *S. aureus* and provides a useful readout of extracellular activity linked to bacterial physiology and virulence-associated regulation (55). Strains carrying pEPSA5-*teg16* exhibited reduced hemolysis compared to vector control strains, as evidenced by decreased zones of clearing surrounding colonies on blood agar (Fig. 4). The representative images shown were obtained under non-inducing conditions, and similar qualitative reductions in hemolysis were observed following xylose induction. This phenotype was consistently observed across independent experiments and across multiple assay formats, including both streak plating and spot-based assays. Time-course analysis further demonstrated that reduced hemolysis in pEPSA5-*teg16* strains was evident at both 24 and 48 hours post-incubation. Together, these results indicate that the *teg16* expression construct is associated with reduced hemolytic activity under the conditions tested.

### Teg16 transiently reduces *codY* expression

Because Teg16 overexpression altered both SigB-associated pigmentation and hemolytic activity, we next asked whether Teg16 also influences broader regulatory pathways associated with metabolism and virulence. We focused on *codY* because prior work identified Teg16 as highly responsive to CodY-dependent regulation, with *teg16* expression reported to increase more than 100-fold in *codY* mutant strains (53). This observation raised the possibility that Teg16 may be embedded within a broader CodY-associated regulatory network. Such a relationship would be consistent with the broader organization of bacterial regulatory systems, in which small RNAs and global transcriptional regulators frequently participate in feedback or feed-forward regulatory circuits. Quantitative RT-PCR analysis revealed that *codY* transcript levels were reduced approximately 5-fold at 30 minutes following induction of Teg16 expression relative to vector control (Fig. 5A). In contrast, no significant change in *codY* expression was observed at the 180-minute time point. This transient reduction in *codY* expression parallels the early regulatory effects observed for rsbV, *sigB*, *asp23*, and *crtM*, suggesting that Teg16 overexpression is associated with early remodeling of regulatory networks beyond the SigB pathway. However, because *codY* repression was not sustained at 180 minutes, these data suggest that Teg16-dependent effects on *codY* are temporally restricted and may reflect an indirect or secondary regulatory response. Together, these findings suggest that Teg16 influences regulatory pathways beyond SigB and may participate in a reciprocal regulatory relationship with the global metabolic regulator CodY. A model summarizing these regulatory relationships is presented in Figure 6.

**Figure 5.**
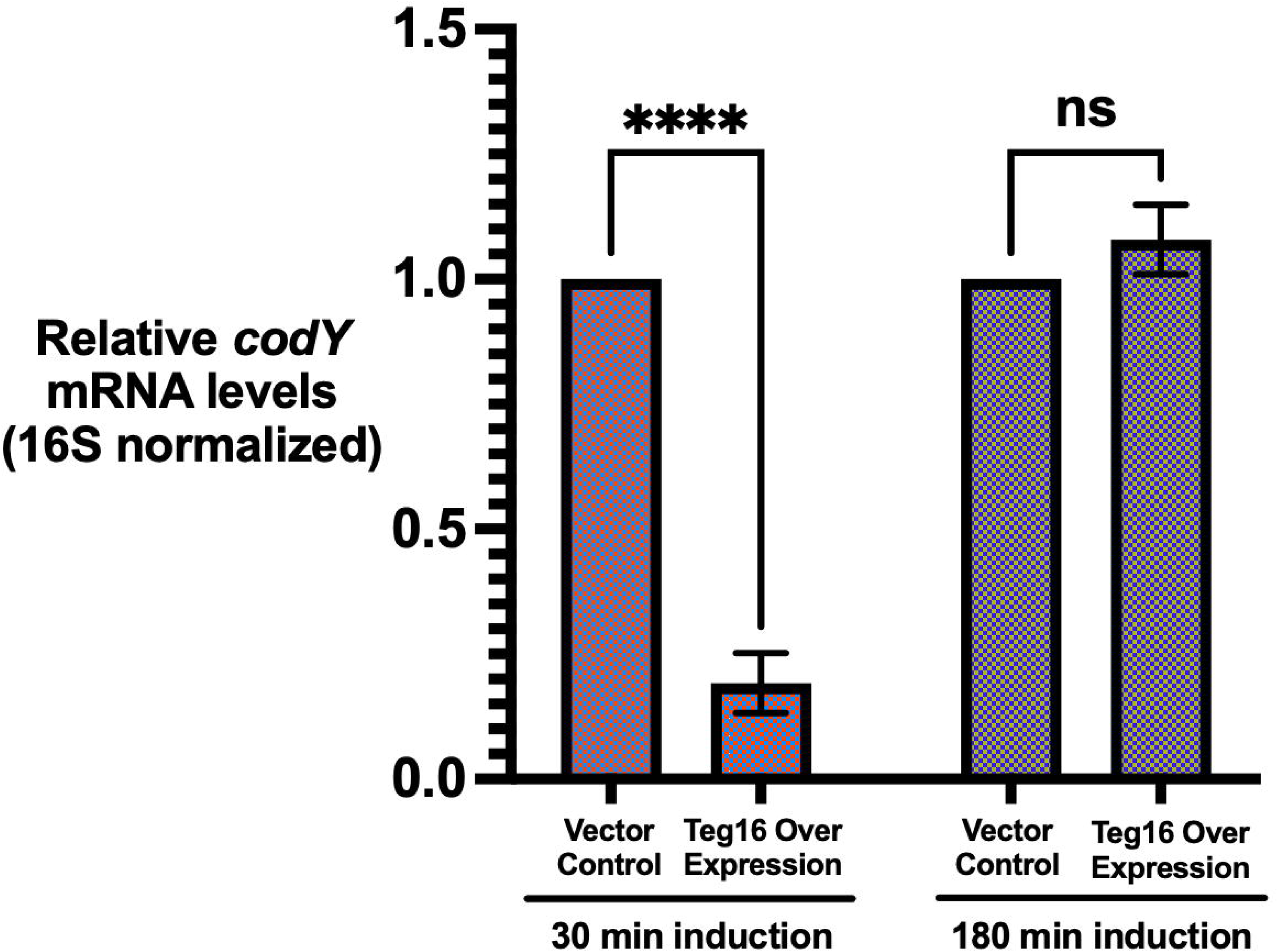
Teg16 represses *codY* expression. To assess whether teg16 influences additional regulatory pathways, *codY* transcript levels were measured following *teg16* overexpression. Quantitative RT-PCR analysis of *codY* mRNA levels was performed using cultures grown to mid-exponential phase (OD_600_: 0.3-0.5) and induced with 2% xylose for 30 or 180 minutes prior to RNA isolation. Transcript levels were normalized to 16S rRNA and expressed relative to vector control. Data represent mean ± SEM from at least three independent biological replicates. Statistical significance was determined using two-way ANOVA followed by Šidák’s multiple comparisons test (\*\*\*\**P* < 0.0001).

**Figure 6.**
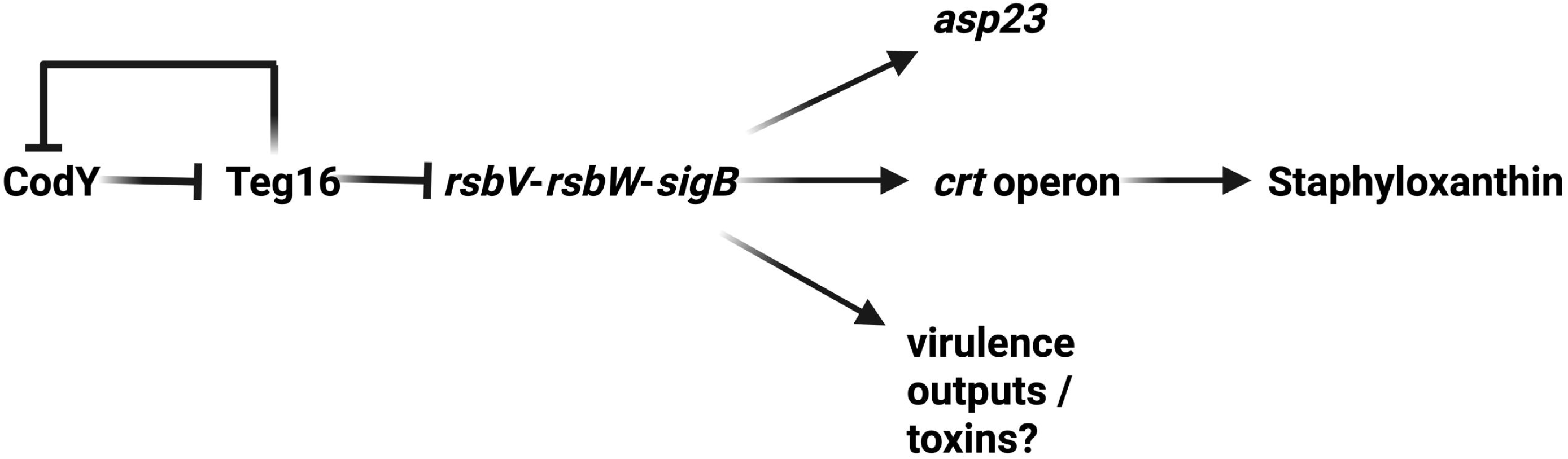
Model for Teg16-mediated regulation of *rsbV*, SigB-dependent outputs, and CodY in *Staphylococcus aureus*. Schematic representation of Teg16 regulatory activity. Teg16 is proposed to repress *rsbV* expression through base-pairing interactions within the translational initiation region, leading to decreased activity of the SigB regulatory pathway and reduced expression of SigB-dependent genes, including those involved in stress response and pigment biosynthesis. Prior studies showed that *teg16* expression is highly responsive to loss of CodY. Our findings suggest that Teg16 may also influence *codY* transcript levels, raising the possibility of a feedback relationship between Teg16 and CodY. The mechanism underlying this interaction remains to be defined. Solid lines indicate experimentally supported regulatory effects, while dashed lines indicate proposed or indirect relationships.

## Discussion

The results presented here identify the small RNA Teg16 as a regulator of *rsbV* expression and downstream SigB-dependent outputs in *Staphylococcus aureus*. Using computational target prediction together with inducible expression analysis, we found that Teg16 overexpression reduces *rsbV* transcript levels and is associated with reduced expression of *sigB* and SigB-dependent genes. These transcriptional changes were accompanied by altered phenotypes, including reduced staphyloxanthin production and decreased hemolytic activity. Together, these findings place Teg16 within the regulatory architecture controlling the SigB stress response pathway. Although the present study relies on inducible overexpression, the consistency across predicted target interaction, transcriptional responses, and phenotypic outputs supports a biologically meaningful role for Teg16 in modulating this pathway. A model summarizing these regulatory relationships is presented in Fig. 6.

The predicted base-pairing interaction between Teg16 and the translational initiation region of *rsbV* provides a plausible mechanism for the observed repression of *rsbV* expression. The location of the predicted interaction, overlapping sequences near the ribosome binding site and start codon, is consistent with a model in which Teg16 interferes with translation initiation and/or promotes transcript destabilization. Following Teg16 induction, *rsbV* transcript levels were reduced at both early and later time points, with the strongest repression observed at 30 minutes. Because *rsbV*, *rsbW*, and *sigB* are organized within the same regulatory operon, reduced *rsbV* expression may have consequences for the broader SigB pathway. Consistent with this model, Teg16 overexpression also reduced *sigB* transcript levels and decreased expression of SigB-dependent genes. The strong reduction in *asp23*, a well-established marker of SigB activity, further supports the conclusion that Teg16 negatively influences SigB-dependent transcriptional output.

The phenotypic changes observed in Teg16-overexpressing strains provide additional support for this regulatory model. Staphyloxanthin production was reduced following Teg16 overexpression, consistent with decreased expression of *crtM* and reduced activity of SigB-associated pigmentation pathways. This phenotype is useful because pigment production provides a visible and quantifiable output of the regulatory changes observed at the transcript level. Strains carrying pEPSA5-*teg16* also displayed reduced hemolytic activity, suggesting that Teg16-associated regulatory effects may extend to extracellular activities linked to virulence-regulatory networks. Because the representative hemolysis images shown were obtained without xylose, this phenotype may reflect basal expression from the pEPSA5-*teg16* construct, although similar qualitative effects were also observed under inducing conditions. Hemolysis in *S. aureus* is controlled by multiple intersecting regulators, so this phenotype should not be interpreted as evidence of a single direct pathway. Rather, the combined effects on *rsbV*, *sigB*, SigB-dependent transcripts, pigment production, and hemolysis support a model in which Teg16 shifts regulatory output across multiple physiologically relevant readouts.

The relationship between Teg16 and CodY provides an additional regulatory connection that may be important for understanding how Teg16 is integrated into broader cellular networks. Prior work demonstrated that *teg16* expression is highly responsive to loss of CodY, placing Teg16 within the CodY-associated RNome (53). That observation suggested that Teg16 may be linked to metabolic state or nutrient-responsive regulation, but the downstream function of Teg16 had not been defined. In the present study, Teg16 overexpression transiently reduced *codY* transcript levels at the early time point. These findings raise the possibility that Teg16 and CodY participate in a reciprocal regulatory relationship. However, the mechanism and directionality of this relationship remain unresolved. The transient nature of the *codY* effect suggests that Teg16-dependent changes in *codY* expression may be indirect, condition-dependent, or buffered by additional regulatory mechanisms over time.

These findings add to a growing view that bacterial sRNAs can influence major regulatory pathways rather than acting only as accessory regulators of isolated transcripts. In *S. aureus*, many sRNAs have been identified through transcriptomic and computational studies, but relatively few have been connected to specific targets, regulatory pathways, or phenotypic outputs. Teg16 has been recognized as part of the *S. aureus* sRNA repertoire, but its function has remained largely uncharacterized. The present study addresses that gap by linking Teg16 to the *rsbV–rsbW–sigB* pathway and to SigB-associated phenotypes. This connection is notable because SigB is most often discussed in the context of partner-switching proteins, transcriptional regulation, and environmental stress inputs. Our findings suggest that post-transcriptional regulation may provide an additional layer of control over this central stress-response pathway.

Several important mechanistic questions remain. First, direct validation will be required to determine whether the predicted Teg16-*rsbV* base-pairing interaction is necessary and sufficient for repression. Target-site mutagenesis, compensatory mutations, and reporter-based assays will be particularly useful for distinguishing direct effects on translation from effects on mRNA stability. Second, additional experiments will be needed to determine whether Teg16 primarily regulates *rsbV* or whether *rsbV* is one component of a broader Teg16 regulon. Third, the physiological conditions that induce endogenous Teg16 expression remain to be defined. This is especially important because inducible overexpression establishes regulatory potential but does not fully define native regulatory context. Future studies combining endogenous expression analysis, target validation, and broader transcriptomic approaches will clarify how Teg16 functions within stress-, metabolism-, and virulence-associated regulatory networks.

In summary, this study identifies Teg16 as a previously uncharacterized sRNA that represses *rsbV* expression and modulates downstream SigB-dependent transcriptional and phenotypic outputs in *Staphylococcus aureus*. The study links Teg16 to the *rsbV–rsbW–sigB* pathway, reduced expression of SigB-dependent genes, decreased staphyloxanthin production, altered hemolytic activity, and transient changes in *codY* expression. These findings provide a foundation for future mechanistic studies of Teg16 target recognition and regulatory function. They also extend the known relationship between Teg16 and CodY by suggesting that Teg16 may participate in a broader regulatory circuit connecting stress response and metabolic control. More broadly, this work supports a model in which post-transcriptional regulation contributes to the fine-tuning of central stress-response pathways in *S. aureus*. This framework should help guide future studies of sRNA-mediated regulation in Gram-positive bacterial physiology and pathogenesis.

## Acknowledgments

We thank members of the Thompson Lab for critical review of this manuscript. We also acknowledge the utilization of resources from the newly established University-Wide Genomics Core Facility at Howard University.

## Author Contributions

Conceptualization: S.P., K.H., A.H., E.T., and K.M.T.; Methodology: S.P., K.H., A.H., E.T., S.L. and K.M.T.; Investigation: S.P., K.H., A.H., E.T., S.L., O.M., and K.M.T.; Validation: S.P., K.H., S.L., O.M., and K.M.T.; Formal analysis: S.P. and K.M.T.; Visualization: S.P. and K.M.T.; Resources: K.M.T.; Supervision: K.M.T.; Project administration: K.M.T.; Funding acquisition: K.M.T.; Writing – original draft: S.P. and K.M.T.; Writing – review and editing: S.P., K.H., A.H., E.T., S.L., O.M., and K.M.T.

K.H. cloned *teg16* into pEPSA5 and verified the construct by sequencing. S.L. and O.M. contributed to repeat pigment analysis and qRT-PCR validation experiments. K.H., A.H., and E.T. contributed to conceptual development of the project based on related unpublished observations. All authors reviewed and approved the final manuscript.

